# HECT-type ligases facilitate autoubiquitination and degradation of other ubiquitin ligases to activate plant immunity

**DOI:** 10.64898/2026.08.06.743213

**Authors:** Zhishuo Wang, Robert O. Mason, Heather Grey, Christos Spanos, Beatriz Orosa-Puente, Steven H. Spoel

## Abstract

The ubiquitin-proteasome system (UPS) serves as the primary proteolytic machinery in eukaryotes, governing intracellular protein turnover to maintain proteome homeostasis. In plants, the HECT-type UPL3/4 ubiquitin ligases play vital roles in developmental and immune signaling. After ubiquitination by pathway-specific E3 ligases, substrates are physically relayed to proteasome-associated UPL3/4 ligases for further modification, which is necessary for their proteasome-mediated degradation. In this study, we investigated if the cellular influence of UPL3/4 extends beyond their direct role in substrate degradation. We discovered that UPL3/4 govern the ubiquitination not only of a broad array of immune-related substrates, but also of many UPS components, including E3 ligases. UPL3 physically interacts with PUB22, a pathway-specific U-box E3 ligase that negatively regulates immunity. PUB22 is controlled by a phospho-switch that converts it from an instable autoubiquitinated state to a stable phosphorylated E3 ligase that marks substrates for degradation. Remarkably, UPL3 only interacted with unphosphorylated PUB22 and facilitated its autoubiquitination-mediated degradation, thereby promoting the accumulation of PUB22 substrates. Moreover, the compromised immune phenotypes of *upl3 upl4* mutant plants were largely dependent on PUB22 and its close paralogues. Thus, UPL3/4 control the stability of immune-related substrates not only through direct ubiquitination, but also indirectly by promoting autoubiquitination of PUB22 ligase and its paralogues. Controlling the stability of autoubiquitinating E3 ligases may be a universal mechanism whereby HECT-type ligases and the proteasomes they associated with, orchestrate cellular proteostasis in eukaryotes.

**Significance Statement:** The ubiquitin-proteasome system (UPS) governs intracellular protein turnover to maintain proteome homeostasis in eukaryotes. Proteasome-associate HECT-type ubiquitin ligases play an important role in processing and degrading substrates delivered to the proteasome by pathway-specific E3 ligases. Here, we discover that in plants, HECT-type ligases not only promote the degradation of substrates, they also modify the E3 ligases that target these substrates to the proteasome. Specifically, HECT-type ligases facilitated or expanded the autoubiquitination of immune-suppressive E3 ligases, resulting in their proteasome-mediated degradation and onset of immunity. Our discoveries suggest that during plant immunity, HECT-type ligases and the proteasomes they associate with, control cellular proteostasis by governing the stabilities of both E3 ligases and their substrates.

## Introduction

As the central proteolytic machinery in eukaryotic cells, the ubiquitin-proteasome system (UPS) maintains homeostasis of ~80–90% of cellular proteins, including pivotal regulators such as tumor suppressors, cell-cycle controllers, signaling receptors, and transcription factors (1–3). Protein substrates are targeted to the UPS by a series of ubiquitin-modifying enzymes. E1 enzymes activate ubiquitin and transfer it to an E2 conjugating enzyme. The E2 then pairs with an E3 ligase that selectively binds a target substrate and catalyzes the transfer of ubiquitin from the E2 to a specific lysine residue on the substrate. Efficient targeting of substrates to the proteasome often requires a polyubiquitin chain consisting of at least four ubiquitin molecules, and is sometimes facilitated by specialized ubiquitin ligases that possess chain-elongation activities (4, 5). The polyubiquitinated substrate is ultimately recognized by dedicated ubiquitin receptors of the 26S proteasome, leading to its degradation.

The 26S proteasome, the central protease of the ubiquitin-proteasome system, is a multi-subunit complex comprising a 20S catalytic core particle and a 19S regulatory particle. The proteasome’s function is regulated not only by its core subunits but also by interacting accessory proteins, including a unique set of ubiquitin ligases (6). In yeast for example, the HECT-type ligase HUL5 associates with the proteasome regulatory particle and plays a critical role in proteasomal substrate degradation (7). HUL5 endows the proteasome itself with ubiquitin ligase activity, enabling the remodeling of substrate-anchored ubiquitin chains at the proteasome (8). The *hul5* mutant exhibits incomplete substrate degradation, indicating that HUL5 is crucial for processive proteolysis (9). Similarly, in plants, the HECT-type Ubiquitin Protein Ligase (UPL) family also associate with the proteasome (10). UPLs are involved in a wide range of biological processes in plants, including immune responses and multiple aspects of growth and development (11–14). Notably, UPL3 and its close paralogue UPL4 have recently been shown to target key transcriptional activators (TAs) essential for hormone-responsive transcriptional reprogramming (10, 15, 16). In a remarkable relay mechanism, pathway-specific E3 ligases transfer substrates to proteasome-bound UPL3/4 for further ubiquitin chain modification and processive degradation (10).

In plants, UPS components have diversified and their number dramatically expanded. Indeed, the importance of the UPS in regulating numerous signaling pathways, particularly in immune responses against pathogens, has been widely reported (17–20). Plant immunity is initiated by cell surface-localized immune receptors that recognize extracellular pathogen-derived molecules and activate pattern-triggered immunity (PTI) (21). In Arabidopsis, for example, the cell-surface immune receptor Flagellin-Sensing 2 (FLS2) recognizes the N-terminal fragment of bacterial flagellin to initiate PTI (22). Activation of FLS2 triggers multiple molecular and physiological defense outputs, including kinase signaling, a rapid reactive oxygen species (ROS) burst, elevated cytosolic calcium, callose deposition at the site of infection and stomatal closure (23). Negative feedback mechanisms reliant on ubiquitin signaling play an important role in preventing excessive or untimely PTI. For example, the U-box E3 ligases PUB12 and PUB13 ubiquitinate the FLS2 immune receptor, thereby promoting its degradation and establishing a critical negative feedback mechanism that prevents excessive PTI signaling (24). Other U-box E3 ligases have also been implicated in regulating plant immune perception and signal transduction. PUB22, PUB23, and PUB24 are functionally redundant E3 ligases that suppress plant immunity (25). PUB22 attenuates defense signaling by directly ubiquitinating and degrading the exocyst component Exo70B2, which controls homeostasis of the FLS2 immune receptor at the plasma membrane (26, 27). Accordingly, dysfunction of PUB22/23/24 enhances immune responses, including increased ROS production, prolonged mitogen-activated protein kinases (MAPKs) activation, and elevated immune gene expression (25, 26). Intriguingly, PUB22 activity is regulated by oligomerization of the U-box domain, which induces its self-or autoubiquitination and proteasomal degradation (28). MPK3 interacts with and phosphorylates PUB22 at Thr-62 and Thr-88 within and adjacent to its U-box domain, respectively, which promotes an oligomer-to-monomer switch, inhibits autoubiquitination and stabilizes PUB22 (28). Thus, regulation of PUB ligase stability and activity is indispensable for appropriate regulation of plant immune responses.

In this study, we discover that HECT-type UPL3/4 ligases interact with numerous UPS components, including immune-associated E3 ligases such as PUB22. We show that UPL3/4 preferentially interacts with the unphosphorylated form of PUB22 and amplifies its autoubiquitination and ultimately proteasome-mediated degradation, thereby fine-tuning downstream plant immune responses. Our findings suggest that HECT-type ligases do not only govern the stabilities of typical cellular substrates, they also process autoubiquitinating E3 ligases to amplify their regulatory reach over the cellular proteome.

## Results

### UPL3/4 regulate numerous UPS components to shape the cellular ubiquitome

UPL3 and UPL4 ligases exert global influence on cellular ubiquitination (29). To profile the UPL3/4-regulated ubiquitome, we conducted proteomics on ubiquitinated proteins pulled down from wild-type (WT) plants and the *upl3 upl4* double mutant. A total of 275 ubiquitin-associated proteins exhibited differential levels in the mutant, with 172 significantly enriched and 103 depleted (*SI Appendix*, Fig. S1A and Dataset S1). Analyses of functional enrichment and protein interactions revealed that these differentially ubiquitinated proteins are strongly associated with plant immune responses, redox homeostasis, and translation initiation (*SI Appendix*, Fig. S1B and C), indicating a broad regulatory role for UPL3/4 in immunity. Notably, the predominant enriched classes of proteins included numerous UPS components, such as proteasome subunits, E2 conjugating and E3 ligase enzymes, and ubiquitin receptors and shuttle proteins (*SI Appendix*, Fig. S1 and Dataset S1). Thus, UPL3/4 may be central regulators of UPS components, thereby extending their reach over the cellular ubiquitome.

### UPL3/4 catalyze the ubiquitination and proteasome-mediated turnover of PUB22

Previously, we reported that in yeast two-hydrid assays, UPL3 also interacts with E3 ligases, including PUB ligases (29). Here, we used co-immunoprecipitation (co-IP) assays to confirm interactions with PUB22 and PUB23, and investigated how UPL3/4 control UPS function. YFP-tagged UPL3 isolated from *35S:YFP-UPL3* (in *upl3-4*) plants specifically pulled down recombinant PUB22-FLAG and PUB23-FLAG *in vitro* (Fig. 1A and *SI Appendix*, Fig. S2). This interaction was further confirmed by *in vivo* co-IP (Fig. 1B), demonstrating that UPL3 directly associates with PUB22 and PUB23. The subcellular localization of PUB22 may correlate with its functional output, as autoinhibited PUB22 at the plasma membrane can be targeted by MPK3 for activation, whereas cytoplasmic PUB22 is associated with active substrate ubiquitination (26, 28). Interestingly, bimolecular fluorescence complementation (BiFC) assays indicated that interaction between UPL3 and PUB22 occurs predominantly at the plasma membrane (Fig. 1C), suggesting UPL3 targets the autoinhibited state of PUB22.

**Figure 1.**
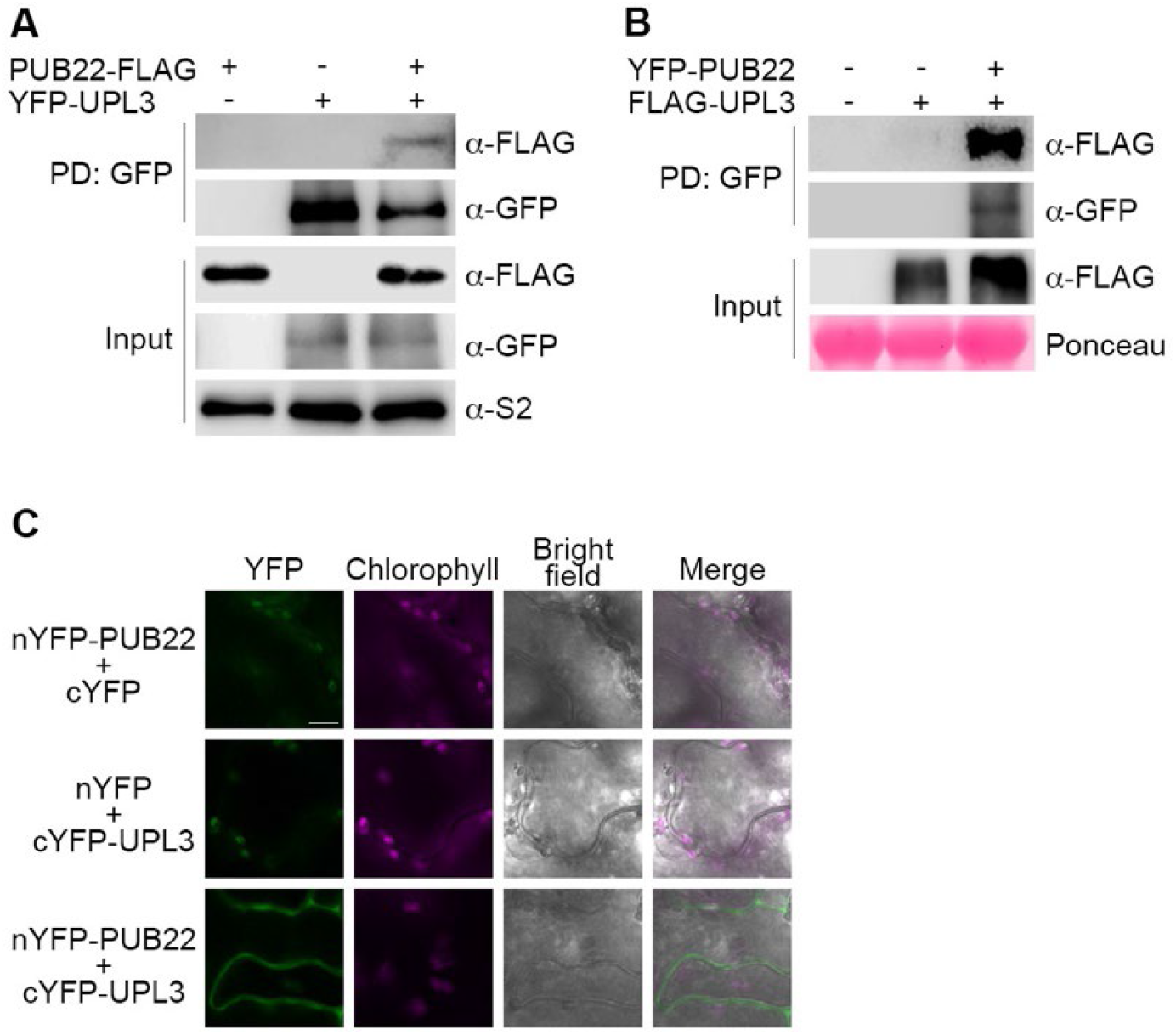
UPL3 physically interacts with PUB22. **(A)** UPL3 interacts with PUB22 *in vitro*. YFP-UPL3 was purified from *35S:YFP-UPL3 (in upl3-4)* plants and incubated with *in vitro* synthesized PUB22-FLAG. Immunoprecipitated proteins were analyzed by immunoblotting with antibodies against GFP and FLAG. The proteasome subunit S2 served as a loading control. **(B)** UPL3 interacts with PUB22 *in vivo*. YFP-PUB22 and FLAG-UPL3 were co-expressed in *N. benthamiana* leaves, protein complexes were isolated and detected by immunoblotting with antibodies against GFP and FLAG. Ponceau S staining is shown as a loading control. **(C)** BiFC analysis of UPL3 and PUB22 interaction in *N. benthamiana*. UPL3 and PUB22 were fused to the C-terminal (cYFP) and N-terminal (nYFP) halves of YFP, respectively. Reconstituted YFP fluorescence and chlorophyll autofluorescence were imaged 3 days post-infiltration. Scale bar = 10 μm.

PUB22 is a negative regulator of PTI and its protein accumulation is tightly controlled through proteasomal degradation (28). Accordingly, treatment with the bacterial PAMP flg22 markedly increased GFP-PUB22 protein abundance, and even higher levels accumulated when proteasome activity was inhibited with MG132 (*SI Appendix*, Fig. S3). Moreover, treatment with the HECT-type ligase inhibitor, heclin, also induced GFP-PUB22 accumulation (*SI Appendix*, Fig. S3) (30), suggesting that HECT-type ubiquitin ligases regulate PUB22 abundance. To examine if turnover of PUB22 is regulated by UPL3 and its close homolog UPL4, we constitutively expressed *YFP-PUB22* in WT plant and *upl3 upl4* double mutants (*SI Appendix*, Fig. S4). Under steady-state conditions, *upl3 upl4* mutants accumulated higher levels of YFP-PUB22 than WT plants (Fig. 2A). This difference was further enhanced upon flg22 treatment, suggesting that UPL3/4 are required for the turnover of PUB22 during immune activation. Furthermore, upon treatment with flg22 and the protein synthesis inhibitor cycloheximide (CHX), YFP-PUB22 was rapidly turned over in the WT background (Fig. 2B and C). By contrast, YFP-PUB22 degradation was significantly impaired in the *upl3 upl4* mutant.

**Figure 2.**
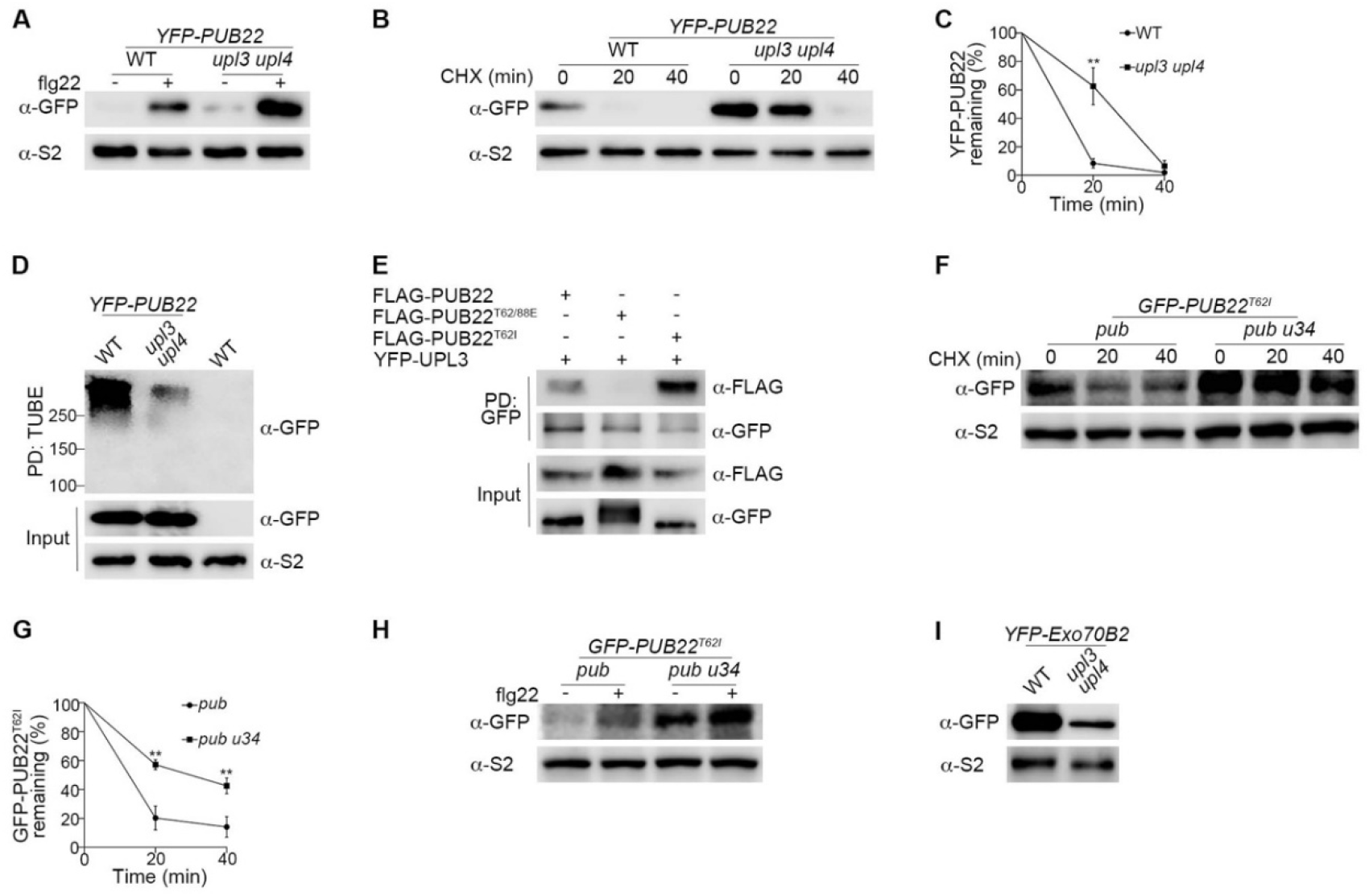
UPL3/4 catalyze polyubiquitination and proteasomal degradation of PUB22. **(A)** The *upl3 upl4* mutant accumulates YFP-PUB22 in response to flg22. Seedlings expressing YFP-PUB22 in the indicated backgrounds were treated with or without 1 μM flg22 for 2 h. YFP-PUB22 was detected by immunoblotting with antibody against GFP. **(B and C)** YFP-PUB22 is stabilized in the *upl3 upl4* mutant background. Seedlings expressing YFP-PUB22 in the indicated backgrounds were treated with 1 μM flg22 for 2 h, then co-treated with 1 μM flg22 and 100 μM CHX. Samples were collected at the indicated time points. In (A) YFP-PUB22 was detected by immunoblotting with antibody against GFP, and proteasome subunit S2 served as a loading control. In (B) YFP-PUB22 levels were quantified by normalizing to S2. Data are mean ± SD (two-tailed *t* test, **P ≤ 0.01; n = 3). **(D)** UPL3/4 mediate polyubiquitination of PUB22. Seedlings were treated with 1 μM flg22 for 2 h, then co-treated with 1 μM flg22 and 100 μM MG132 for 4 h. Ubiquitinated proteins were enriched with GST-TUBE and ubiquitinated YFP-PUB22 was detected by immunoblotting with antibody against GFP. **(E)** UPL3 preferentially interacts with unphosphorylated PUB22. YFP-UPL3 was transiently co-expressed with FLAG-PUB22, FLAG-PUB22^T62/88E^, or FLAG-PUB22^T62I^ in *N. benthamiana*. Protein complexes were isolated and detected by immunoblotting against GFP and FLAG. **(F and G)** Loss of *UPL3/4* function stabilizes GFP-PUB22^T62I^. Seedlings expressing YFP-PUB22^T62I^ were treated as in (B). YFP-PUB22^T62I^ was detected by immunoblotting against GFP, and S2 served as a loading control (F). Protein levels were quantified as in (C). Data are mean ± SD (two-tailed *t* test, **P ≤ 0.01; n = 3) (G). **(H)** UPL3/4 regulate the accumulation of YFP-PUB22^T62I^. Seedlings expressing YFP-PUB22^T62I^ were treated with or without 1 μM flg22 for 2 h. YFP-PUB22 was detected by immunoblotting against GFP. **(I)** flg22-induced YFP-Exo70B2 accumulation is impaired in the *upl3 upl4* mutant. Plants carrying *pUBQ10:YFP-Exo70B2* were treated with flg22 for 1 h, and protein levels were analyzed by immunoblotting against GFP and S2.

To determine if UPL3/4 mediate ubiquitination of PUB22, we isolated total cellular ubiquitinated proteins from flg22-treated plants expressing *YFP-PUB22* using tandem ubiquitin-binding entities (TUBEs) (31), and subsequently probed for the presence of YFP-PUB22. Compared to WT, ubiquitination of YFP-PUB22 was largely compromised in the *upl3 upl4* mutant, as evidenced by a substantial reduction in high-molecular-weight, polyubiquitinated species (Figure 2D). Thus, UPL3/4 catalyze the polyubiquitination and subsequent proteasome-mediated degradation of PUB22.

### Autoubiquitination of PUB22 is a prerequisite for further ubiquitination and degradation via UPL3/4 ligases

A low level of ubiquitinated YFP-PUB22 was still present in the *upl3 upl4* mutant (Figure 2D), which we reasoned may be due to PUB22 autoubiquitination (28). Therefore, we examined the possibility that PUB22 autoubiquitination is a necessary step for its recognition and subsequent polyubiquitination by UPL3/4. Given that PUB22 autoubiquitination occurs *in trans* via oligomerization of its U-box domain (28), we investigated if UPL3/4 target oligomeric PUB22 for degradation. Mutant *upl3 upl4* plants accumulated more oligomeric PUB22 than WT, especially after flg22 treatment (*SI Appendix*, Fig. S5A), suggesting UPL3/4 can target the oligomeric self-ubiquitinating state of PUB22. To test this further, we constitutively expressed a catalytically inactive GFP-PUB22^W40A^ variant that largely lacks ubiquitin ligase activity (25). Unlike the wild-type GFP-PUB22 protein (Figure 2D), mutation of *UPL3* and *UPL4* did not reduce further the residual ubiquitination of GFP-PUB22^W40A^ (*SI Appendix*, Fig. S5B), indicating autoubiquitination of PUB22 is a prerequisite for subsequent ubiquitination by UPL3/4.

Since phosphorylation of PUB22 by MPK3 blocks its oligomerization and thereby reduces autoubiquitination (28), we co-expressed UPL3 with a phosphonull PUB22 variant (GFP-PUB22^T62I^) that due to enhanced hydrophobic interactions exhibits enhanced oligomerization, and a phosphomimetic PUB22 variant (GFP-PUB22^T62/88E^) that is predominantly monomeric (28). Co-IP assays demonstrated UPL3 strongly interacted with the oligomeric, phosphonull GFP-PUB22^T62I^ variant, whereas its interaction with the monomeric, phosphomimetic GFP-PUB22^T62/88E^ variant was barely detectable (Fig. 2E). Next, we generated transgenic Arabidopsis plants constitutively expressing *GFP-PUB22*^*T62I*^ to confirm UPL3/4 target the phosphonull variant for degradation (*SI Appendix*, Fig. S6A). Because of functional redundancy between PUB22 and its paralogs PUB23 and PUB24 (25), we expressed *GFP-PUB22*^*T62I*^ in a *pub22 pub23 pub24 upl3 upl4* quintuple mutant background. Indeed, after CHX treatment, degradation of the flg22-induced phosphonull PUB22^T62I^ variant was impaired when both *UPL3* and *UPL4* were dysfunctional (Fig. 2F and G). Furthermore, while GFP-PUB22^T62I^ was stabilized by flg22 treatment in the *pub22 pub23 pub24* triple mutant, its levels were already markedly higher in untreated *pub22 pub23 pub24 upl3 upl4* quintuple mutants and did not rise much further upon flg22 treatment (Fig. 2H). These data demonstrate that UPL3/4 only target the autoubiquitination-competent, oligomeric form of PUB22 and facilitate its degradation.

Next, we examined if UPL3/4-mediated degradation of PUB22 also affects the stability of PUB22’s own substrates. As an E3 ligase itself, PUB22 ubiquitinates and targets Exo70B2 for proteasome-mediated and endovacuolar degradation pathways, constituting an important negative feedback loop to repress plant immune responses (26, 32). Compared to WT plants, *upl3 upl4* mutants exhibited elevated levels of PUB22 (Fig 2A-2C) but accumulated dramatically lower levels of YFP-Exo70B2 (Fig. 2I and *SI Appendix*, Fig. S6B). Thus, UPL3/4 not only regulate the stability of PUB22, but also of its downstream immune-associated substrates.

### UPL3/4 activate PTI responses by targeting PUBs

A major hallmark of immune responses is the rapid production of ROS triggered by the detection of pathogen-derived PAMPs during the early stages of infection (33). PUB22/23/24 ligases are negative regulators of plant immunity and dampen ROS production (25, 26). Because UPL3 and UPL4 are also major regulators of plant immunity and target PUB22/23 for degradation (10, 29), we investigated if PAMP-induced ROS production is compromised in *upl3* and *upl4* mutants. Compared to WT plants, the *upl3* single mutant exhibited a modest reduction in flg22-induced ROS production, whereas no significant difference was observed in the *upl4* mutant (Fig. 3A and B). By contrast, both the amplitude and total accumulation of the flg22-induced ROS burst were markedly reduced in the *upl3 upl4* double mutant (Fig. 3A and B). Thus, both UPL3 and UPL4 are required for the full activation of ROS production upon pathogen perception. Next, we examined if reduced ROS accumulation in *upl3 upl4* mutants may be a consequence of failing to degrade PUB22. To test this we utilized the *pub22 pub23 pub24* triple mutant to remove functional redundancy between PUB paralogs (25). Consistent with previous reports (26), the triple mutant exhibited enhanced PTI responses, as evidenced by elevated ROS production (Fig. 3C and D). Remarkably, however, mutation of *PUB22/23/24* in the *upl3 upl4* double mutant background restored ROS levels to near WT levels (Fig. 3C and D). Although the quintuple mutant continued to display a prolonged ROS burst similar to that of the *pub22 pub23 pub24* mutant (Fig. 3C), these data demonstrate that UPL3/4 enhance the ROS burst by controlling the levels of PUB22 and its paralogs.

**Figure 3.**
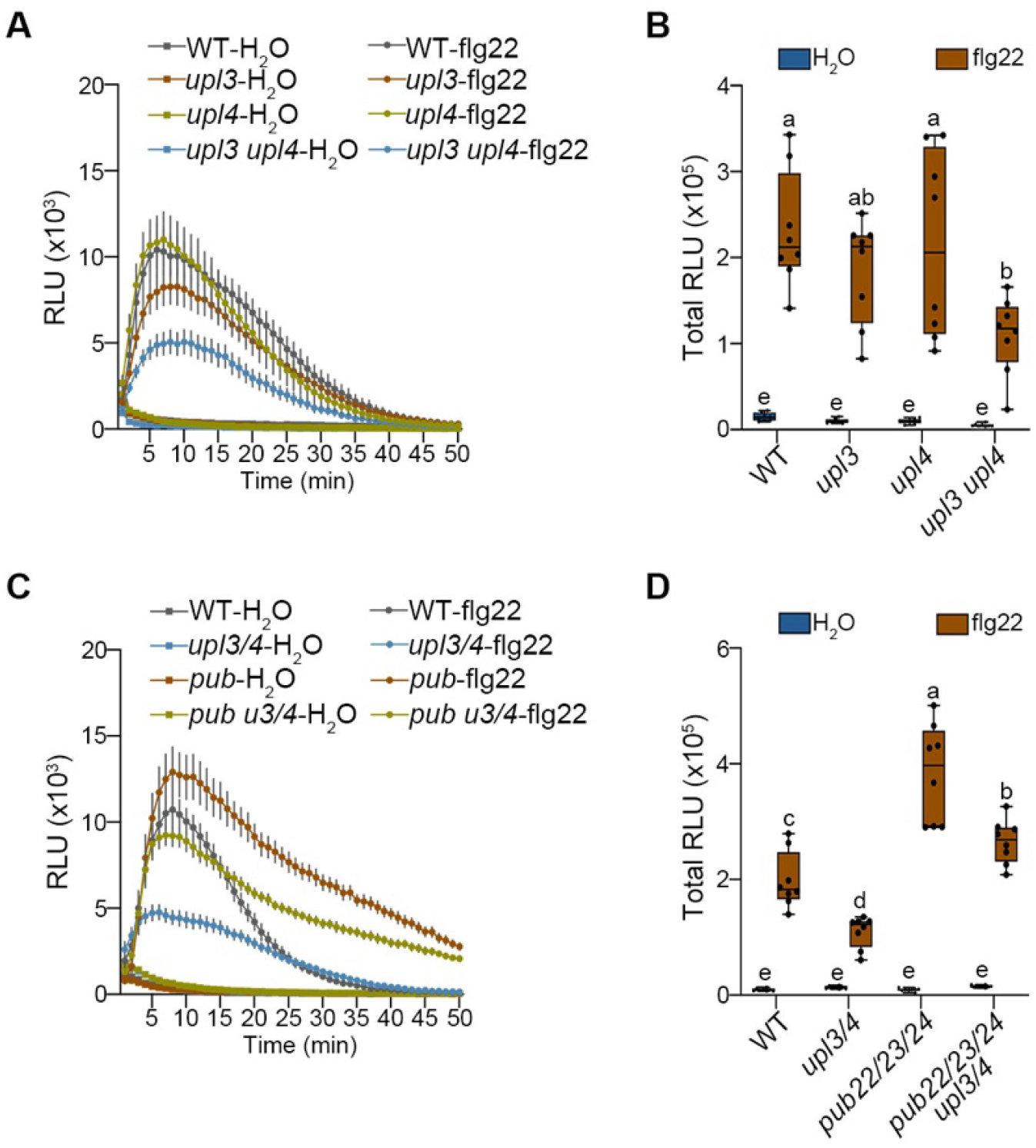
UPL3/4 enhance the flg22-induced ROS burst by targeting PUB22. **(A and B)** UPL3 and UPL4 are required for ROS production in response to flg22. Leaf discs from 4-week-old plants were treated with 1 μM flg22. In (A) ROS production is shown as relative light units (RLU) and data are presented as the mean ± SE (n = 8). In (B) total ROS production was measured over a 50 min period. Data are presented as the mean ± SD, lowercase letters indicate significant differences between samples (Tukey HSD ANOVA test; α = 0.05, n = 8). **(C and D)** Mutations of PUB22/23/24 (i.e. *pub*) restores ROS production in the *upl3 upl4* mutant. Measurements were performed as in (A) and (B).

Due to the energy trade-off between defense and growth, sustained activation of PTI is associated with attenuated plant growth (34). We therefore tested if the UPL3/4-PUB22/23/24 regulatory module impacts this trade-off in response to PAMP perception. Flg22 treatment strongly inhibited root growth in WT plants, whereas *upl3* and *upl4* single mutants exhibited reduced sensitivity (Fig. 4A and B). Notably, this phenotype was further enhanced in the *upl3 upl4* double mutant (Fig. 4A and B). We then asked if the reduced flg22 sensitivity in *upl3 upl4* mutants results from a failure to degrade the negative regulators PUB22/23/24. As expected, the *pub22 pub23 pub24* triple mutant displayed enhanced PTI responses, as evidenced by a magnified inhibition of root elongation (Fig. 4C and D). Intriguingly, introducing *PUB* mutations into the *upl3 upl4* background largely phenocopied the *pub22 pub23 pub24* triple mutant phenotype (Fig. 4C and D). Taken together, these findings indicate that UPL3/4-mediated ubiquitination and degradation of the PUB22/23/24 ligases are essential for establishing rapid as well as sustained PTI responses.

**Figure 4.**
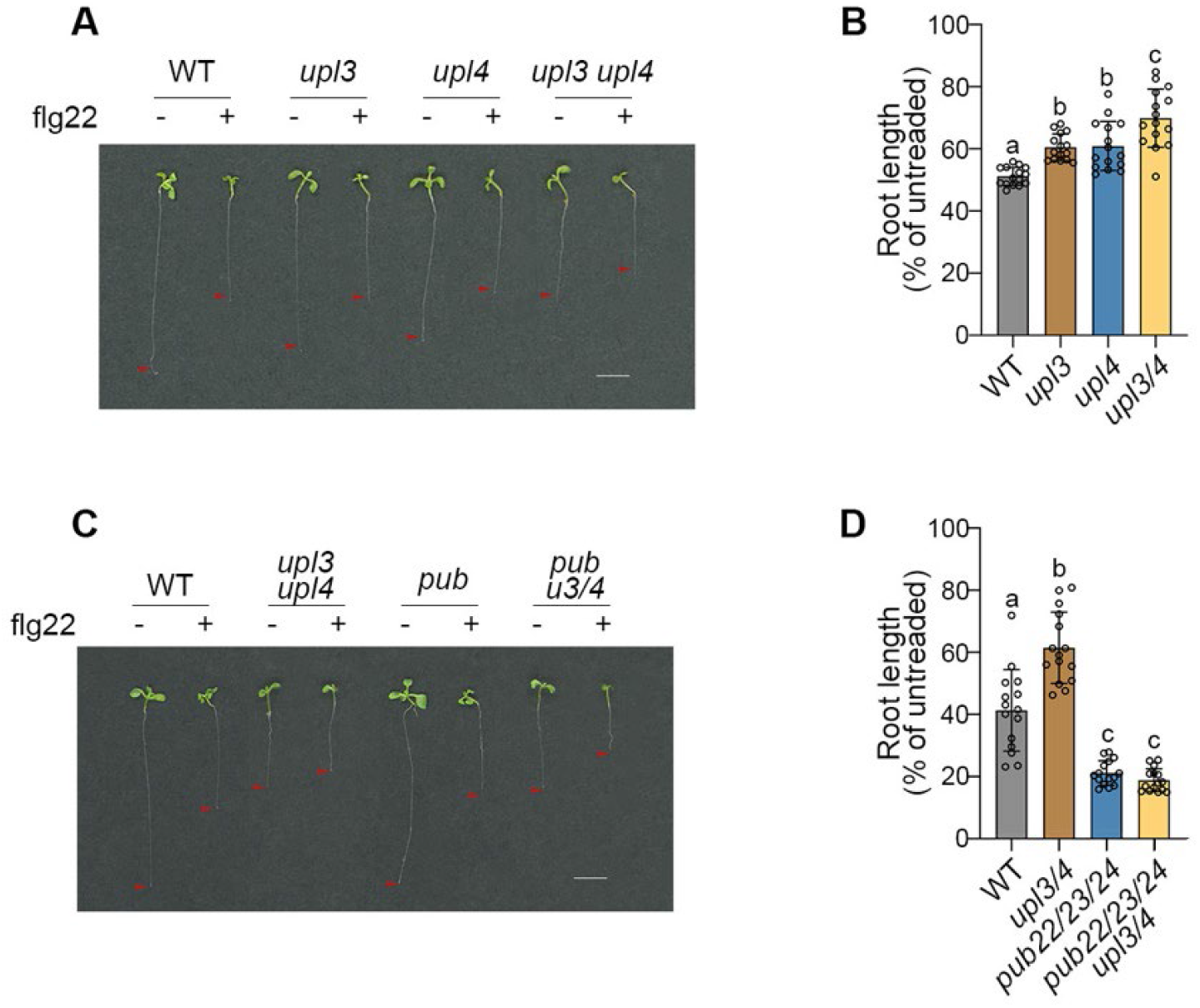
UPL3/4 are required for sustained PUB22-mediated PTI responses. **(A and C)** Four-day-old seedlings were transferred to MS agar medium supplemented with or without 1 μM flg22. Representative morphological phenotypes were analyzed 6 days post-transfer. Arrowheads indicate primary root tips. Scale bar = 50 mm. **(B and D)** Primary root growth following flg22 treatment, presented as a percentage of the untreated control root length. Data points represent the mean ± SD (n = 15). Lowercase letters indicate statistically significant difference between samples (Tukey HSD ANOVA test; α = 0.05).

### UPL3/4-mediated degradation of PUB22/23/24 regulates plant immunity

Because flg22-triggered responses were attenuated in the *upl3 upl4* mutant, we next examined if UPL3/4 are essential for PTI against a bacterial leaf pathogen. When spray-inoculated with *Pseudomonas syringae pv. tomato* (*Pst*) DC3000, the *upl3* mutant displayed enhanced disease susceptibility, whereas the *upl4* mutant exhibited resistance levels similar to that of WT plants (Fig. 5A and B). Given the functional redundancy of UPL3 and UPL4 in PTI, inoculation of the *upl3 upl4* double mutant resulted in severely enhanced *Pst* DC3000 growth, exceeding the level observed in the *upl3* single mutant (Fig. 5A and B). Strikingly, disease resistance was restored in the *upl3 upl4 pub22 pub23 pub24* quintuple mutant to levels comparable to WT (Fig. 5C and D), suggesting that overaccumulation of PUB paralogs is chiefly responsible for the enhanced disease susceptible phenotype of the *upl3 upl4* double mutant. Hence, our collective data reveal that UPL3/4-mediated ubiquitination and proteasomal degradation of PUB22/23/24 are essential for establishing effective immunity.

**Figure 5.**
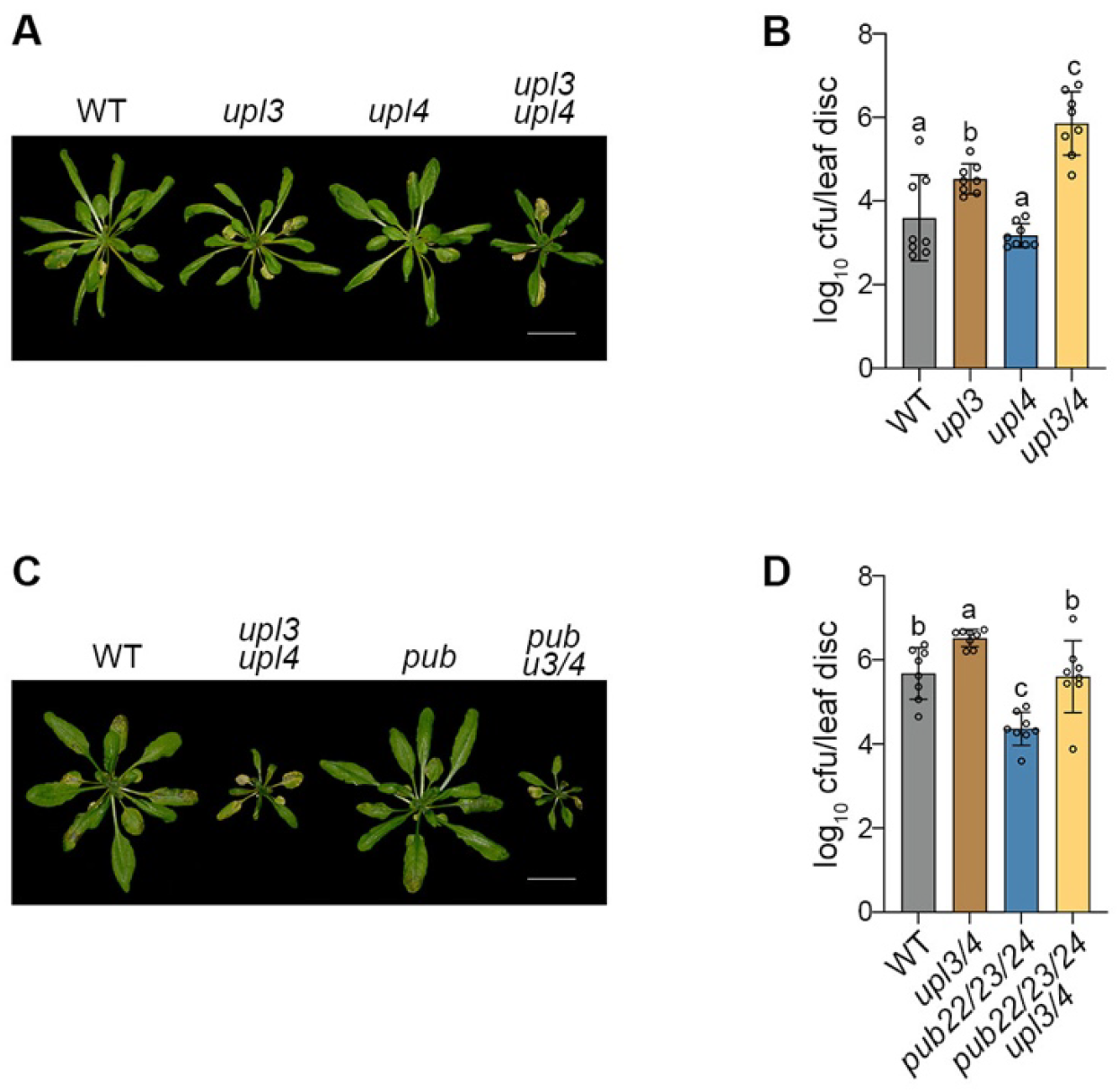
UPL3/4-mediated regulation of PUB22/23/24 activity is required for immunity. (A and C) Disease symptoms in adult plants at 4 days post spray inoculation (dpi) with a *Pst* DC3000 suspension of 5 x 10^8^ cfu/mL. Scale bar = 1 cm. (B and D) Bacterial growth was determined and data points represent the mean ± 95% confidence limits. Lowercase letters indicate significant differences between samples (Tukey HSD ANOVA test; α = 0.05, n = 8).

## Discussion

Protein ubiquitination is a major mechanism for regulating cellular homeostasis in eukaryotes by targeting substrates for proteasomal degradation. Recent evidence from multiple organisms demonstrates that proteasome-associated HECT ligases catalyze substrate polyubiquitination at the proteasome, a ubiquitin chain remodeling step essential for processive substrate degradation (8, 35). Here, we demonstrate that the HECT-type ligases UPL3 and UPL4 not only target a broad array of immune-related proteasome substrates, but also the E3 ligases that govern the fate of these substrates. We propose that by controlling the stability of self-ubiquitinating E3 ligases, such as PUB22/23/24, HECT-type ligases and the proteasomes they associate with, drastically extend their influence over the cellular proteome to regulate plant immunity.

In Arabidopsis, seven HECT-type ubiquitin ligases have been characterized and are implicated in diverse biological processes (12, 29, 36). Our previous work demonstrated that UPL3/4 are required for immune responses induced by salicylic acid (SA), the central hormone that activates local and systemic immunity (29, 37). SA-induced transcriptional reprogramming is modulated by the UPS, which dynamically fine-tunes the transcriptional activity of NPR1, a key regulator of SA signaling (38). Notably, UPL3/4-mediated polyubiquitination is essential for clearing inactive NPR1 from SA-responsive promoters, thereby allowing re-initiation of the transcription cycle (10). Accordingly, loss of *UPL3* and *UPL4* function impairs SA responses, including reduced induction of immune-related genes and abolished SA-induced pathogen resistance (29). Consequently, UPL ligases have emerged as prime targets for pathogens seeking to disable plant immunity. An effector from the potato cyst nematode, *Globodera pallida*, interacts with the potato functional homolog of Arabidopsis UPL3, which is essential for activating nematode-induced transcriptional responses (39). Likewise, *Pst* DC3000 promotes virulence by secreting the effector HopM1 to target proteasome-associated UPLs and suppress host proteasome function (40).

Our findings demonstrate that UPL3/4 modulate not only SA-mediated immunity, but also target many UPS components, including E3 ligases in early immune signalling (Fig. S1). Here, we show that UPL3/4 fine tune PTI by ensuring the degradation of autoubiquitinated PUB22, a negative regulator of PTI responses (Figs. 1 and 2). Compared to WT plants, the *upl3 upl4* mutant displayed a significantly reduced ROS burst in response to flg22, which correlated with decreased trade-offs between growth and immunity, and enhanced susceptibility to pathogens (Figs. 3, 4 and 5). The phenotypes of *upl3 upl4* mutants were largely rescued by introducing mutations in *PUB22/23/24*. Notably, while disease resistance was restored to WT levels in the *pub22 pub23 pub24 upl3 upl4* quintuple mutant, it did not phenocopy the enhanced disease resistance of the *pub22 pub23 pub24* triple mutant (Fig. 5C and D). This implies that defects in SA-mediated local and systemic immunity also contribute to the compromised resistance of the *upl3 upl4* mutant as reported previously (10, 29, 37). Collectively, these data establish UPL3/4 as essential regulators that restrict pathogen invasion through multi-layered control of the plant immune system.

Like UPL3/4, several HECT-type E3 ligases have been reported to ubiquitinate other E3 ligases. For example, in human cells, the HECT-type ligase UBE3A (E6AP) targets the RING E3 ligase Ring1B (RNF2) for proteasomal degradation, thereby reducing histone H2A monoubiquitination and modulating transcriptional repression (41, 42). Moreover, both NEDD4 and ITCH HECT-type ligases ubiquitinate the RING-type E3 ligase CBL, although the physiological consequences of this modification remain poorly understood (43). Interestingly, in these cases Ring1B and CBL ligases also exhibit self-ubiquitination, but this activity is distinct from their ubiquitination by HECT-type ligases and in case of Ring1b, even has opposing effects on its stability. Here, we demonstrated that in plants, self-ubiquitination of PUB22 ligases is a prerequisite for further ubiquitination by HECT-type UPL3/4 ligases and degradation by the proteasome (Fig 2). These findings suggest that self-ubiquitination is not always sufficient to target E3 ligases for proteasomal degradation and instead, require further modification by HECT-type ligases. Curiously, many E3 ligases associate with the proteasome, which may be to relay their ubiquitinated substrates directly to the proteasome (10). HECT-type ligases may help the proteasome distinguish between E3 ligases that deliver their ubiquitinated cargo versus E3 ligases that are themselves destined for degradation. In case of PUB22, its UPL3/4-mediated ubiquitination and subsequent proteasome-mediated degradation are tightly linked to a phosphorylation event that determines its oligomer-to-monomer switching. Oligomerization is a well-characterized regulatory mechanism for U-box E3 ligases, and in several cases, oligomer formation is essential for catalytic activity. For example, structural studies of the yeast U-box ligase Prp19 show that dimerization via the U-box is essential for its function, as its disruption causes cell death (44). Similarly, the U-box E3 ligase CHIP requires dimerization for activity, despite only one U-box domain in the dimer being catalytically exposed and functional (45, 46). Our results show that UPL3/4 preferentially interact with and promote the degradation of unphosphorylated, oligomeric, and self-ubiquitinated PUB22 (Fig. 2 E to H), the effects of which extended to the stability of PUB22’s own substrates (Fig 2I). Thus, HECT-type ligases interpret extensive interplay between phosphorylation, protein conformation and ubiquitination to determine the stabilities of E3 ligases and their cellular substrates. Because some PUB22 substrates, such as Exo70B2, are degraded through endovacuolar pathways (32), UPL3/4 may amplify their reach over the proteome not only through proteasomal pathways, but also via other ubiquitin-mediated degradation pathways.

In summary, our findings uncover a multi-layered regulatory mechanism for ubiquitin chain assembly on E3 ligases, enabling precise control of their stability. This mechanism relies on a two-step relay, in which ubiquitin chains are first initiated on the E3 ligase via autoubiquitination and subsequent further ubiquitination by proteasome-associated HECT-type ligases promotes their processive degradation. This relay of autoubiquitinated E3s to proteasome-associated HECT ligases may govern the stability of a complete network of E3 ligases as well as their repertoires of cellular substrates, thereby ensuring spatiotemporal regulation of downstream cellular signaling events, including immunity in plants.

## Materials and Methods

### Plant materials and growth conditions

All Arabidopsis plant materials used in this study are of the Col-0 ecotype. The *upl3* and *upl4* single, *upl3 upl4* double, and *pub22 pub23 pub24* triple mutants, and the complemented line *35S:YFP-UPL3 (upl3-4)* have been previously described (10, 25). For experiments on adult plants, Arabidopsis seeds were stratified at 4°C for 2 days prior to sowing on soil. Plants were grown in a chamber maintained at 21°C under a 16-h-light/8-h-dark photoperiod with a light intensity of 100 μmol.m^−2^.s^−1^. Day and night humidity were set to 65% and 55%, respectively. For experiments on seedlings, seeds were surface-sterilized with 100% ethanol for 5 minutes with rotation, then with 10% bleach for 5 min with rotation, and finally rinsed three times with sterile distilled water. Seeds were then plated on Murashige and Skoog (MS) agar medium and incubated under the same environmental conditions described above.

### Protein-protein interactions

For *in vitro* pull-down assays, YFP-UPL3 was purified from *35S:YFP-UPL3 (upl3-4)* plants using GFP-Trap Agarose beads (ChromoTek) according to the manufacturer’s instructions. The bead-bound proteins were then incubated with 500 µL of interaction buffer (10 mM Tris-Cl [pH 7.5], 150 mM NaCl, 0.5 mM EDTA, protease inhibitors cocktail) containing FLAG-tagged PUB22 protein, which was produced using a cell-free synthesis system (47), for 1 h at 4°C. Agarose beads were washed 3 times with interaction buffer, and bound proteins were eluted by boiling at 80°C for 10 min in 1× SDS sample buffer containing 50 mM DTT.

For *in vivo* co-IP assay in *Nicotiana benthamiana, Agrobacterium tumifaciens* cultures carrying the indicated constructs were harvested and resuspended in an infiltration buffer containing 10 mM MgCl_2_ and 10 μL/L 6-benzyladenine. Suspensions were infiltrated into leaves of *N. benthamiana* and harvested 3 days later. YFP-tagged proteins were purified using GFP-Trap Agarose beads (ChromoTek) and eluted as described above. Eluted proteins were separated by SDS-PAGE, and interacting proteins were detected by immunoblotting with antibodies specified in the figure legends.

For the BiFC assays, the split-YFP system was employed, constructs were transiently expressed in *N. benthamiana* leaves via *A. tumefaciens*-mediated infiltration (48). Samples were collected 3 days post-infiltration and imaged using a Leica SP8 confocal microscope.

### Protein analysis

For protein extraction, frozen tissue was homogenized in extraction buffer containing 50 mM Tris-HCl (pH 7.5), 150 mM NaCl, 5 mM EDTA, 0.1% Triton X-100, 0.2% Nonidet P-40, and protease inhibitor cocktail (50 μg/mL N-p-Tosyl-L-phenylalanine chloromethyl ketone (TPCK), 50 μg/mL Nα-Tosyl-L-lysine chloromethyl ketone hydrochloride (TLCK), 0.6 mM phenylmethylsulfonyl fluoride (PMSF)), unless otherwise indicated. Protein extracts were mixed with 1× SDS sample buffer, with or without 50 mM DTT, incubated at 80°C for 10 min, and then separated by SDS-PAGE. To monitor protein degradation, 2-week-old seedlings were pretreated with 1 µM flg22 for 2 h and then co-treated with 1 µM flg22 and 100 µM CHX. Samples were collected at the indicated time points and proteins extracted as previously described.

To analyze protein polyubiquitination, 2-week-old seedlings were treated with 1 µM flg22 for 2 h, followed by co-treatment with 1 µM flg22 and 100 µM MG132 for an additional 4 h. Polyubiquitinated proteins were subsequently enriched using glutathione-S-transferase (GST)-tagged TUBE (tandem ubiquitin binding entity) as described previously (38). Ubiquitinated YFP-PUB22 was detected by immunoblotting with an anti-GFP antibody.

### LC-MS/MS analysis

Leaf tissue was homogenized in an extraction buffer containing 1× PBS, 0.05% Triton, 0.05% Nonidet P-40, 10 mM NEM, 100µM MG132, 200µM PPI3, and protease inhibitor cocktail. The cell lysate was incubated with a recombinant HaloTag-Ubiquilin fusion protein (MRC PPU, University of Dundee) to enrich ubiquitinated proteins. Protein complexes were then isolated using Magne® HaloTag® Beads (Promega).

Beads were resuspended in 50 µL of 8 M urea in 50 mM ammonium bicarbonate (ABC). Proteins were reduced with 100 mM DTT for 30 min at room temperature and alkylated with 100 mM chlorodoacetamide in the dark for 30 min. Digestion was initiated with LysC (enzyme-to-protein ratio 1:50, w/w) for 4 h at room temperature. The urea concentration was then diluted below 2 M by adding 50 mM ABC, followed by overnight trypsin digestion (1:50, w/w) with shaking. The digest was acidified with 0.1% trifluoroacetic acid (TFA) and desalted using StageTips (49). Peptides were eluted with 80% acetonitrile (ACN) in 0.1% TFA, concentrated to 1 µL by vacuum centrifugation, and adjusted to 5 µL with 0.1% TFA for LC-MS/MS analysis.

### MS Data Analysis

Raw mass spectrometry files were processed using MaxQuant (version 1.6.1.0) with Andromeda search engine (50, 51). Spectra were searched against the *Arabidopsis thaliana* reference proteome from UniProt (release August 2019). For the first search, peptide tolerance was set to 20 ppm, while for the main search it was set to 4.5 ppm. Isotope mass tolerance was 2 ppm and maximum charge set to 7. Digestion mode was set to specific with trypsin allowing maximum of two missed cleavages. Carbamidomethylation of cysteine was set as a fixed modification. Oxidation of methionine, and the diGly residue on lysine were set as variable modifications. Absolute protein quantification was performed as previously described (52). Peptide and protein identifications were filtered to a 1% false discovery rate (FDR).

### Phenotypic analysis

Reactive oxygen species (ROS) were measured as previously described (53), with minor modifications. Leaf discs from 4-week-old plants were floated on water in a 96-well plate overnight. The following day, an elicitation solution containing 5 μg/mL Horseradish Peroxidase (HRP) and 0.2 μM Luminol, with or without 1 μM flg22, was added to each well. Luminescence was measured at 1 min intervals using an Infinite M200 PRO Plate Reader (Tecan) with a 0.3 s signal integration time.

For the root growth inhibition assay, seeds were germinated and grown on MS agar medium for 4 days. Seedlings were then transferred to fresh MS medium supplementing with or without 1 μM flg22. Primary root length was measured 6 days after transfer using ImageJ software.

For *Pst* DC3000 inoculation, bacteria cells were collected from overnight cultures and resuspended in ddH_2_O containing 0.04% Silwet L-77 to a final OD600 of 1.0. Four-week-old plants were spray-inoculated with bacterial suspension. Leaves were harvested at 4 days post-inoculation (dpi), and surface-sterilized in 70% ethanol for 30 s, rinsed in ddH_2_O for 30 s, and homogenized in 10 mM MgCl_2_. The ground leaf suspension was serially diluted and plated on LB agar plates and incubated at 30°C for 2 days to determine colony-forming units (CFU).

### Gene expression measurements

Total RNA extraction, cDNA synthesis, and quantitative polymerase chain reaction (qPCR) were performed as described (10). Expression levels of the GFP fusion constructs (forward 5’-AAGCTGACCCTGAAGTTCATCTGC-3’, reverse 5’-CTTGTAGTTGCCGTCGTCCTTGAA-3’) were analyzed using gene-specific primers and normalized to *UBQ5* (forward 5’-CCAAGCCGAAGAAGATCAAG-3’, reverse 5’-ACTCCTTCCTCAAACGCTGA-3’).

### Data, Materials, and Software Availability

Mass spectrometry data have been deposited to the PRIDE repository with the dataset identifier PXD072854. All data are available in the main text or the supplementary materials.

## Acknowledgments

We thank Prof. Marco Trujillo (University of Hamburg) for kindly providing genetic resources and research materials related to PUB ligases. We thank the MRC PPU Reagents and Services facility (MRC PPU, College of Life Sciences, University of Dundee, Scotland, mrcppureagents.dundee.ac.uk) for providing the HaloTag-Ubiquilin expression plasmid. This work was supported by the European Research Council (ERC) under the European Union’s Horizon 2020 research and innovation program, grant agreement no. 101001137 (to S.H.S.) and no. 101126022 (to B.O.-P.), the Biotechnology and Biological Sciences Research Council (BBSRC) grant BB/S016767/1 (to S.H.S.), and by funding for the Wellcome Discovery Research Platform for Hidden Cell Biology [226791], and we gratefully acknowledge support from the Proteomics core.

