## Supplemental Information for "HECT-type ligases facilitate autoubiquitination and degradation of other ubiquitin ligases to activate plant immunity"

**Zhishuo Wang[a], Robert O. Mason[a], Heather Grey[a], Christos Spanos[b], Beatriz Orosa-Puente[a,c,*], Steven H. Spoel[a,*]**
[a]Institute of Molecular Plant Sciences, School of Biological Sciences, University of Edinburgh, Edinburgh, UK.
[b]Institute of Cell Biology, School of Biological Sciences, University of Edinburgh, Edinburgh, UK.
[c]Centro Singular de Investigación en Química Biolóxica e Materiais Moleculares (CiQUS), Universidade de Santiago de Compostela, 15782 Santiago de Compostela, Spain.

[*]Corresponding authors: Beatriz Orosa-Puente and Steven H. Spoel

**This PDF file includes:**

Figures S1 to S6
Legend for Datasets S1

**Other supporting materials for this manuscript include the following:**

Datasets S1

### Figures

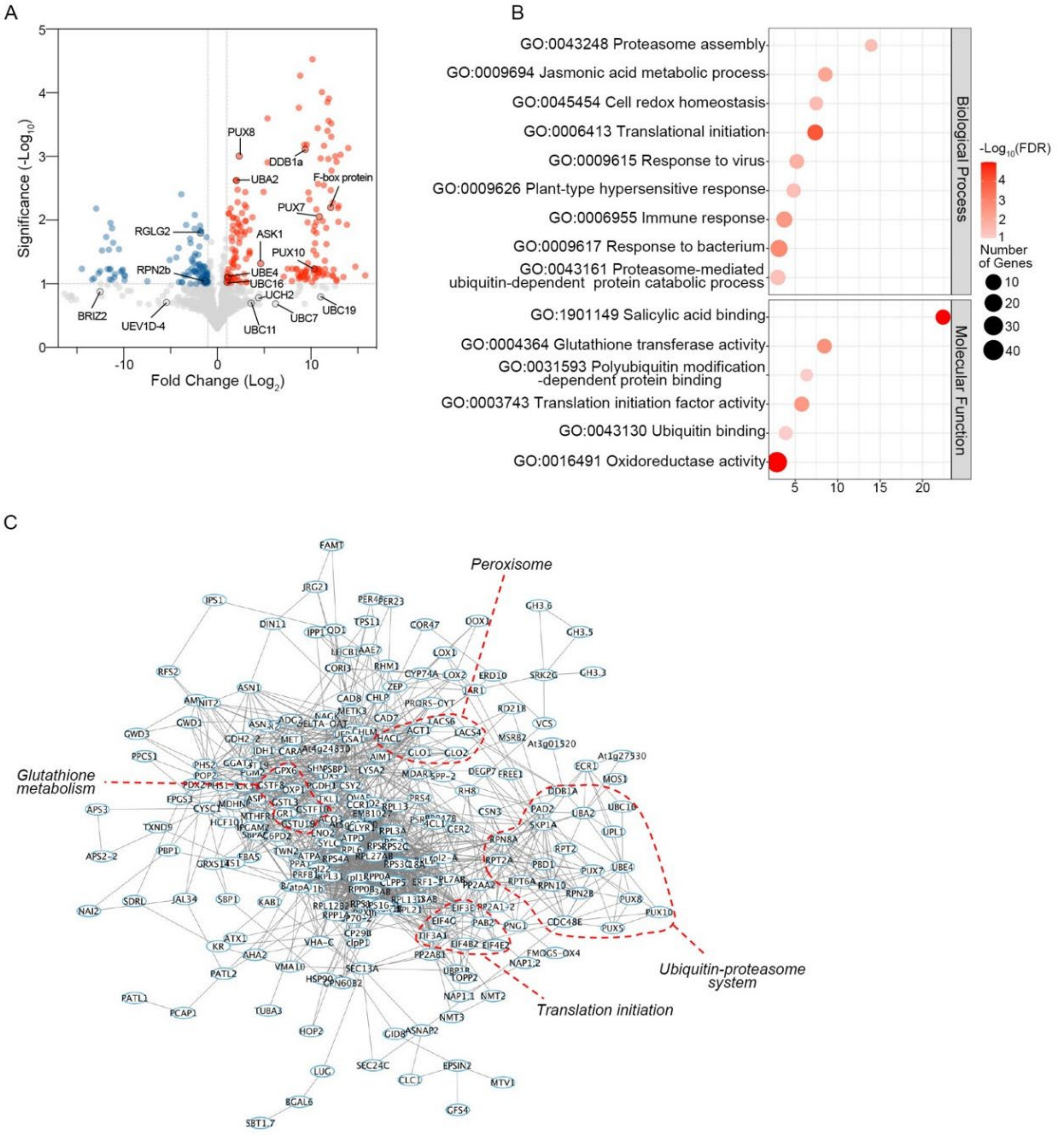


**Fig. S1. Proteomic profiling of the UPL3/4-dependent ubiquitome.**

**(A)** Volcano plot of ubiquitinated proteins enriched (red) or depleted (blue) in the *upl3 upl4* mutant compared to WT. Dashed lines indicate significance thresholds ($p \leq 0.1$, FC $\geq 2$). **(B)** GO enrichment analysis of differentially ubiquitinated proteins. Analysis was performed using ShinyGO V0.74 Gene Ontology Enrichment Analysis tool (http://bioinformatics.sdstate.edu/go/) with default parameters. **(C)** Interaction network of enriched proteins was generated with STRING and subsequently analyzed and visualized using Cytoscape.

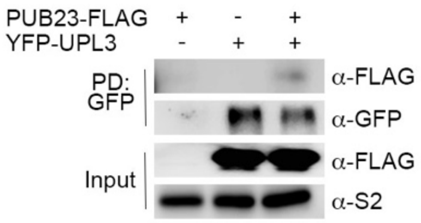


**Fig. S2. UPL3 interacts with PUB23 *in vitro*.**

Purified YFP-UPL3 from *35S:YFP-UPL3 (in upl3-4)* plants was incubated with *in vitro* synthesized PUB23-FLAG. Co-immunoprecipitated proteins were detected by immunoblotting with anti-GFP and anti-FLAG antibodies. The proteasome subunit S2 served as the loading control.or paste legend here. Paste figure above the legend.

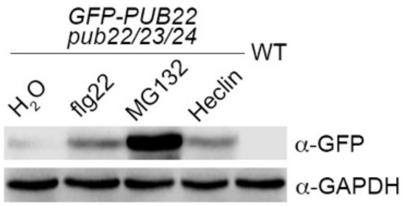


**Fig. S3. Heclin treatment induces PUB22 accumulation.**
Seedlings were treated for 2 h with 1 µM flg22, 0.1 mM MG132, or 0.2 mM Heclin. Protein accumulation was analyzed by immunoblotting against GFP and GAPDH antibody.

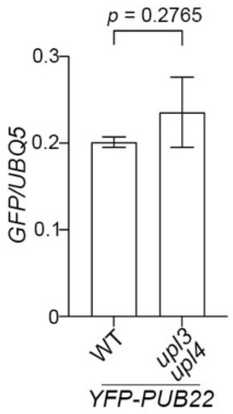


**Fig. S4. Expression of transgenic *YFP-PUB22* in WT and *upl3 upl4* mutants.**

*YFP-PUB22* transgene expression was normalized to the constitutively expressed *UBQ5*. Data represent the mean ± SD. Significant differences were assessed using Student's *t*-test, and *p* value is shown (n=3).

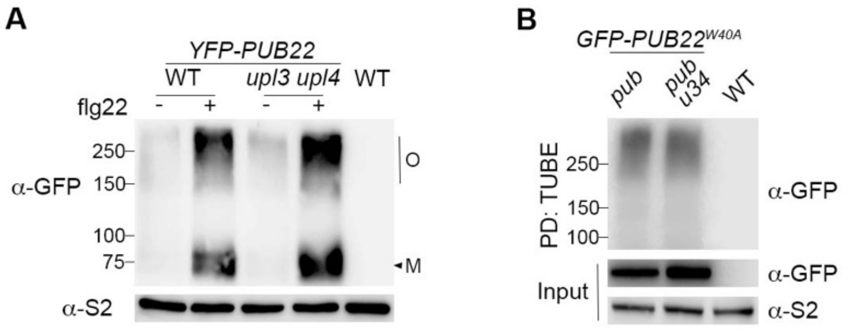


**Fig. S5. Accumulation of PUB22 oligomers and ubiquitinated GFP-PUB$^{W40A}$ in mutants lacking functional *UPL3/4*.**

**(A)** Flg22-induced PUB22 oligomers accumulate in the *upl3 upl4* mutant. Protein extracts were incubated with SDS sample buffer without DTT, and analyzed by immunoblotting with antibody against GFP (O, oligomer; M, monomer). Protein extracts with DTT were probed for S2 as a loading control. **(B)** Autoubiquitination of PUB22 is required for UPL3/4-mediated polyubiquitination. Plants expressing GFP-PUB22$^{W40A}$ in the indicated backgrounds were co-treated with 1 μM flg22 and 100 μM MG132 for 4 h. Ubiquitinated GFP-PUB22$^{W40A}$ was detected by immunoblotting with antibody against GFP.

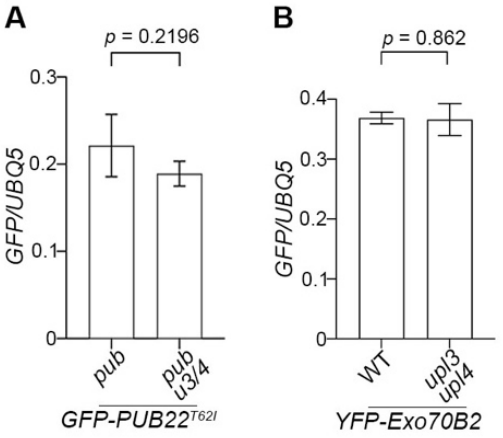


**Fig. S6. Expression of *GFP-PUB22$^{T62I}$* (A) and *YFP-Exo70B2* transgenes in WT and *upl3 upl4* mutants.**

*GFP-PUB22$^{T62I}$* **(A)** and *YFP-Exo70B2* **(B)** transgene expression was normalized to constitutively expressed *UBQ5*. Data represent the mean ± SD. Significant differences were assessed using Student's *t*-test, and *p* value is indicated (n=3).

**Dataset S1 (separate file). Differential ubiquitin-associated proteins in WT versus *upl3 upl4* double mutant plants.**

Proteomics was conducted on ubiquitinated proteins pulled down with a recombinant HaloTag-Ubiquilin fusion protein from wild-type (WT) plants and the *upl3 upl4* double mutant. For each genotype 3 biological repeats were included (R1-R3). This identified 275 ubiquitin-associated proteins with differential levels in the mutant versus WT, with 172 significantly enriched and 103 depleted ($p \leq 0.1$, FC $\geq 2$).
